# Rapid Quantification of First and Second Phase Insulin Secretion Dynamics using In vitro Platform for Improving Insulin Therapy

**DOI:** 10.1101/2023.04.03.535481

**Authors:** Sikha Thoduvayil, Jonathan S. Weerakkody, Mackenzie Topper, Manindra Bera, Jeff Coleman, Xia Li, Malayalam Mariappan, Sathish Ramakrishnan

## Abstract

High-throughput quantification of the first- and second-phase insulin secretion dynamics is intractable with current methods. The fact that independent secretion phases play distinct roles in metabolism necessitates partitioning them separately and performing high-throughput compound screening to target them individually. We developed an insulin-nanoluc luciferase reporter system to dissect the molecular and cellular pathways involved in the separate phases of insulin secretion. We validated this method through genetic studies, including knockdown and overexpression, as well as small-molecule screening and their effects on insulin secretion. Furthermore, we demonstrated that the results of this method are well correlated with those of single-vesicle exocytosis experiments conducted on live cells. Thus, we can quantitatively determine the number of vesicles that fuse when a stimulus is applied. We have developed a robust methodology for screening small molecules and cellular pathways that target specific phases of insulin secretion, resulting in a better understanding of insulin secretion, which in turn will result in a more effective insulin therapy through the stimulation of endogenous glucose-stimulated insulin secretion.

## 1. Introduction

Understanding insulin secretion from pancreatic beta cells holds great promise for solving diabetic pathology and developing therapeutic approaches to prevent its onset.^1-5^ The administration of insulin is necessary for lowering blood glucose levels in individuals with diabetes who cannot produce sufficient insulin on their own. Insulin therapy presents several challenges, including the possibility of hypoglycemia and weight gain.^6^ Therefore, improving endogenous insulin secretion is an important goal, but it requires a deeper understanding of insulin secretion in order to accomplish this objective. Insulin secretion is mediated by major intracellular signals (Calcium influx, hormones, ATP, cAMP, and phospholipid-derived molecules such as diacylglycerol (DAG) and inositol 1,4,5-trisphosphate (IP3)).^7-10^ Through a better understanding of insulin secretion, we may be able to identify potential intervention targets to improve insulin secretion in patients with diabetes.

Insulin secretion occurs in two phases: the first phase occurs within ten minutes following the administration of glucose and the second phase, occurs over a prolonged period, lasting up to 40 minutes.^11-16^ While the physiological and metabolic significance of the biphasic response to a beta cell is not yet fully understood, the available data indicates that the first and second phases play distinct roles in their regulation of insulin secretion. A good first-phase insulin secretion is essential for maintaining glucose homeostasis, as it inhibits hepatic glucose production and postprandial hyperglycemia.^17,18^ The second phase of insulin release, for instance, controls beta-cell mass^19^ and plays a strong role in promoting muscle glucose uptake.^18,20^ It is essential to develop a high-throughput assay that addresses each phase of insulin secretion independently in order to understand the molecular mechanism behind biphasic secretion and how it results in diabetic complications.

Generally, insulin secretion is measured using ELISAs, which require expensive antibodies and multi-step liquid handling.^21,22^ As an alternative, assays involving time-resolved measurements (TR) with fluorescence resonance energy transfer (FRET) have become commercially available at a slightly lower cost, but advanced software and precise liquid handling steps make these assay less accessible.^23,24^ Previous attempts at measuring insulin secretion, by fusing GFP or luciferase to the end of the proinsulin peptide, have failed because of protein misfolding or due to them being stuck in the endoplasmic reticulum.^25^ The development of a smaller luminescent protein, nanoluc (Nluc) (19.1 kDa, monomeric), through directed evolution from luciferases of the deep-sea shrimp (Oplophorus gracilirostris),^26^ have demonstrated the ability of Nluc to produce strong luminescence upon catalyzing the furimazine substrate (half-life > 2 h).^27,28^ This propels them as an ideal co-agonist pair for high-throughput screening and an important biological tool for protein-protein interaction studies *in vitro*.

Here, we report a high-throughput method for assessing first and second-phase insulin secretion dynamics in beta cells by measuring Nluc secretion along with insulin upon glucose stimulation. To validate this system, we have tested known secretagogues and suppressors, Ca^2+^ signaling, gene overexpression and knockdown, and pathological disease models to visualize their impacts on the first and second phases of exocytosis. This will set a path forward to rapidly perform small molecule compound screens to identify potentiators and inhibitors to target a cellular signaling pathway.

## 2. Materials and Methods

### 2.1. Cell culture

The clonal beta-cell line 832/13, derived from the parental INS-1 rat insulinoma cells, was purchased from Millipore Sigma. 832/13 cells were cultured at 37°C and 5% CO_2_ in complete RPMI medium (RPMI-1640 (Sigma Cat. No. R0883) supplemented with 10% FBS (Cat. No. ES-009-B), 10 mM HEPES (Cat. No. TMS-003-C), 2 mM L-Glutamine (Cat. No. TMS-002-C),1 mM sodium pyruvate (Cat. No. TMS-005-B), and 0.05 mM beta-mercaptoethanol (Cat. No. ES-007-E).

### 2.2 Generation of Proinsulin-Nluc stable cell line

INS-1 cells were seeded in 60 mm dish and transfected with 4.5 μg pCI-insulin-Nluc and 0.5 μg pcDNA3.1/Zeo empty vector. Cells stably expressing Insulin-Nluc were selected by treating with 150 μg/ml Zeocin (Invitrogen, Lot# 1937479) for 3 weeks. The individual clones were manually picked, and the expression of Insulin-Nluc was analyzed by luciferase assay.

### 2.3 Transfection Studies

INS-1 cells were transiently transfected with proinsulin-Nluc, proinsulin-eGFP, and syntaxin4/SNAP23 DNA constructs. Proinsulin-NanoLuc in pLX304 was a gift from David Altshuler (Addgene plasmid # 62057; http://n2t.net/addgene:62057 ; RRID:Addgene_62057)^29^ For transient transfection, 2 × 10^5^ INS-1 cells were plated for 6 well plates a day prior to the experiment. Then, 2.5 μg of DNA was mixed with Lipofectamine reagent 3000 (Invitrogen, cat no 2489212) and followed manufacturer protocol suitable for 6 well plates. After a 6 h incubation (37 °C, 5% CO_2_), Opti-MEM medium was replaced with fully supplemented with complete RPMI medium. After 24 hours of transfection, the cells were then washed and used for vesicle isolation and experimental studies.

### 2.4 Small Interfering RNA (siRNA) Transfection

Cells were seeded with complete RPMI medium onto six-well tissue culture plates. After reaching 80% confluence, the cells were transfected with VAMP8 siRNA (Product no. NM_031827, #SASI_Rn01_00112721, Millipore Sigma, USA) at a final concentration of 10 nM using Lipofectamine RNAiMAX transfection reagent according to the manufacturer’s protocol. After transfection for 48 h, cells were treated with 18 mM glucose and 0.25 mM IBMX for luciferase assay. For a single vesicle fusion assay in live cells, we transfected proinsulin-pHlourin DNA along with a siRNA transfection.

### 2.4 Immunofluorescence Assay

The cells were grown on a Poly-D-Lysine Coated MatTek glass bottom dish for immunofluorescence images. The cells were rinsed in PBS prior to being fixed in 4% paraformaldehyde for 30 minutes at room temperature. The fixed cells were washed with PBS and permeabilized with 0.25% Triton X-100 for 15 minutes. After 1 h of blocking with 2% bovine serum albumin, cells were incubated with the anti-Insulin antibodies at room temperature for 2 h. The cells were washed with PBS and incubated with the secondary antibodies conjugated with either 488 nm or 647 nm for 1 h at room temperature. Finally, the cell nuclei were labeled with Hoechst 33342 nucleic acid stain for 10 min at room temperature and the immunofluorescence images were acquired using a Leica Confocal SP8 fluorescence microscope.

### 2.5 Western Blot Analysis

Cells were scratched from the plate and lyzed them directly with electrophoresis (Laemmli) sample buffer and boiled the mixture at 95–100 °C for 5 min. Proteins were separated by SDS/PAGE method, and performed western blots based on the Biorad western blot protocol. The blots were stained with anti-insulin antibodies (Mouse monoclonal, #66198-1-1g, Proteintech), Syntaxin-1 (rabbit polyclonal, #A4133, Abclonal), VAMP8 (mouse monoclonal, #MA5-32502, Invitrogen) depending on the protein analyzed. The concentration of antibody used in the western blots was 1:1000 and developed with ECL western blot substrate.

### 2.6 Insulin dense core granules isolation

Cells were cultured in three 150 mm plates. After 80 % confluency, the plates were used for the insulin granules isolation. All the steps were performed at 4 °C. First, the plates were washed three times using ice-cold PBS. Remove adherent cells by gently scraping with a plastic cell scraper in 5 mL ice-cold PBS buffer. Pellet the harvested cells by centrifugation for 5 min at 200 × g, 4 °C. Resuspended the pellets in 1ml lysis buffer (32 mM sucrose, 5 mM HEPES,1 mM ATP and protease inhibitor) and performed 20 strokes to lyze the cells using a Dounce homogenizer. Centrifuge the lyzed samples for 5 min at 1000 × g, 4 °C to pellet the nuclear components. The supernatant collected from the previous step was separated on a discontinuous OptiPrep density gradient containing five concentrations (8.8%, 13.2%, 17.6%, 23.4%, and 30%) of 2 mL each. SW41 rotor was used to centrifuge the tubes for 75 minutes at 100,000 x g, 4 °C. Twenty fractions (650 uL each) were individually collected from the top of the OptiPrep gradient.

### 2.6 Insulin secretion assay

INS-1 cells were seeded in 6-well plates with an initial density of 1.0 × 10^6^ cells per well and were grown to confluence. The glucose-stimulated insulin secretion is performed in HBSS (HEPES balanced salt solution): 114 mmol/L NaCl, 4.7 mmol/L KCl, 1.2 mmol/L KH_2_PO_4_,1.16 mmol/L MgSO_4_, 20 mmol/L HEPES, 2.5 mmol/L CaCl_2_, 25.5 mmol/L NaHCO_3_, and 0.2% bovine serum albumin, pH 7.2. Cells were washed twice with HBSS + 2.5 mM glucose and left for 15 mins. After 15 mins, 1 mL of HBSS buffer containing 18 mM glucose and 0.25 mM IBMX diluted is added to each well. Every step was carefully performed to ensure that cells were not detached from the plates. Every 5 mins, 100 μL of the supernatant solution was collected from the same wells to measure a kinetic response. The solutions were loaded into a 96-well white plate with 50 μM furimazine substrate to measure the luminescence. The following equations were employed to calculate the luminescence (RLU) at each time point:

Calculated RLU = (Sample intensity at time (t) × dilution factor) + Σintensity (t-1, t-2, etc.) Dilution factor = (total volume remaining in well, time t) ÷ (collected volume size).

### 2.7 Quantification of Nluc secretion

Cells in the wells were lyzed with Promega cell-lysis buffer to estimate the residual Nluc in the cells. We used luciferase intensity to estimate the amount of Nluc secretion at a given time interval. First-phase insulin secretion was defined as insulin released in the first 10 minutes after the addition of HBSS containing 18 mM glucose and 0.25 mM HBSS, and second-phase insulin release was defined as insulin release after 10 minutes as previously described.

## 3 Results

### 3.1. Design of Nluc high-throughput assay to quantify first and second-phase insulin secretion

We transfected a plasmid encoding proinsulin-Nluc luciferase into INS1 cells, a well-established model cell line for glucose-stimulated insulin secretion (GSIS), cultured in a 6-well tissue culture plate. When vesicle maturation occurs, pH-sensitive prohormone convertases cleave the Nluc luciferase and co-secretes insulin, which can be measured quantitatively and used in high-throughput screenings. (Figure 1A). Nluc luciferase and insulin secretion were measured in response to increasing glucose concentrations, confirming that these cells faithfully secrete Nluc luciferase along with insulin (R^2^ = 0.95, Figure 1B). To determine if the assay could reliably reproduce *in vivo* physiology upon glucose stimulation, we applied 18 mM glucose to the cells expressing the reporter. Luciferase secretion showed prolonged and sustained release over time, as predicted (Figure 1C). We observed that the transfection efficiency was consistent with Lipofectamine 3000 and Opti-Mem medium (Figure S1). To enable an understanding of genes and compounds that influence the first or second phase of secretion particularly, the luminescence assay was adapted in a high-throughput format, and we measured luminescence intensity every 5 mins, similar to the ELISA assays reported in literature. An 18 mM glucose solution was added to each well, and the supernatants were collected every five minutes without disturbing the monolayer of cells. For further investigations, we chose 400-600 nm wavelengths for luminescence intensity acquisition based on the luminescence spectrum generated by purified Nluc protein (Figure S2). We applied the transformation (see methods) to deconvolve the measured luminescence intensity at each time point (Figure 1D). It is also possible to measure luminescence intensity by either directly adding substrate furimazine to each well or by transferring the supernatant to a white 96-well plate before adding furimazine. As expected, we can clearly discern the luciferase secretion at every time interval and are therefore confident we are reproducing physiology, demonstrating that human islets perfusion assays for drug testing *in vivo* are unnecessary, along with expensive ELISA tests. To facilitate high-throughput assays and reduce the need for transfection with plasmids, INS1 cells were engineered to stably express insulin-Nluc, and their morphology, proliferation rate, and viability was compared to those of wild type.

**Figure 1.**
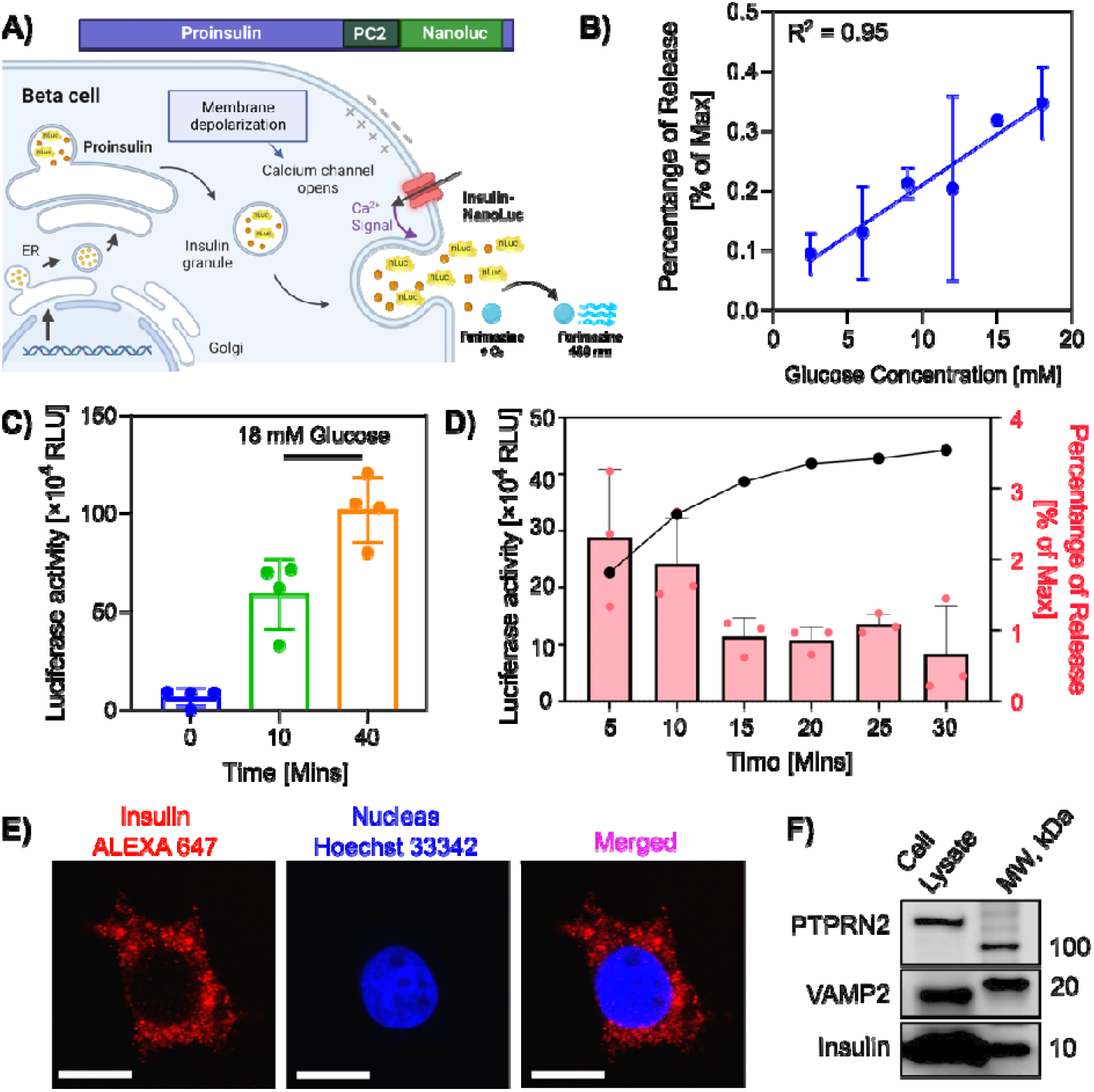
Nanoluc luciferase assay to quantitatively dissect the physiology of insulin biphasic release. **(A)** Scheme representation of creation and expression of Nluc in the cells. An increase in Ca^2+^ influx induced by glucose stimulation results in the co-secretion of Nluc and insulin, which when reacted with furimazine substrate produces high-intensity luminescence. **(B)** The plot shows the correlation between luciferase intensity from INS-1E cells expressing the Nluc vs different concentrations of glucose; data represent mean ± SD (n = 2). **(C)** The representative bar chart shows the luciferase intensity measured at 0, 10 and 40 mins after glucose stimulation; data represent mean ± SEM (n = 4). Cells co-secrete Nluc and insulin for 40 mins, whereas the maximum release happens in the first 10 mins. **(D)** Quantification of Nluc intensity at a defined time point provides information about the probability of release; data represent mean ± SEM (n = 3). **(E)** Representative fluorescence micrographs of cells stained with anti-insulin antibodies tagged with Alexa-647 and Hoechst-33258 (Scale bar: 10 μm)**(F)** Representative western blot image of insulin from cell lysate expressing Nluc.

We independently validated the cells expressing insulin by targeting them with anti-insulin antibodies using an immunofluorescence approach with a confocal microscope (Figure 1E) and western blotting (Figure 1F). We isolated the insulin-containing granules using a density centrifugation method to confirm the presence of Nluc by measuring the luminescence from each fraction (Figure S3). The isolated insulin granules were characterized by western blotting using antibodies against VAMP2 and PTPRN2 (phogrin), which are transmembrane proteins associated with insulin granules (Figure S4). In parallel, insulin-eGFP plasmids were transfected into cells and isolated granules were imaged using a total internal reflection fluorescence microscope (TIRF). The insulin granules containing eGFP fluorescence and Phogrin colocalized perfectly to the vesicles (Figure S5). These findings conclude Nluc was correctly targeted to insulin-containing secretory vesicles via endogenous proinsulin processing, and its activity is preserved.

### 3.2. Validating the assay with known secretagogues and suppressors

As part of our evaluation, we incubated the cells with a panel of known secretagogues and suppressors to determine whether the assay could replicate insulin secretion and biphasic release. The presence of 3-isobutyl-1-methylxanthine (IBMX), which is known to increase cyclic adenosine monophosphate (cAMP) levels, results in increased luciferase secretion in the presence of 18 mM glucose (Figure 2A). This enhanced secretion of IBMX follows the trend of glucose concentrations (Figure 2B). Based on these findings, IBMX enhanced luciferase secretion compared to glucose, making it an important parameter for monitoring the two phases separately. Therefore, all the subsequent studies were carried out in the presence of 0.25 mM IBMX and were considered as a control.

**Figure 2.**
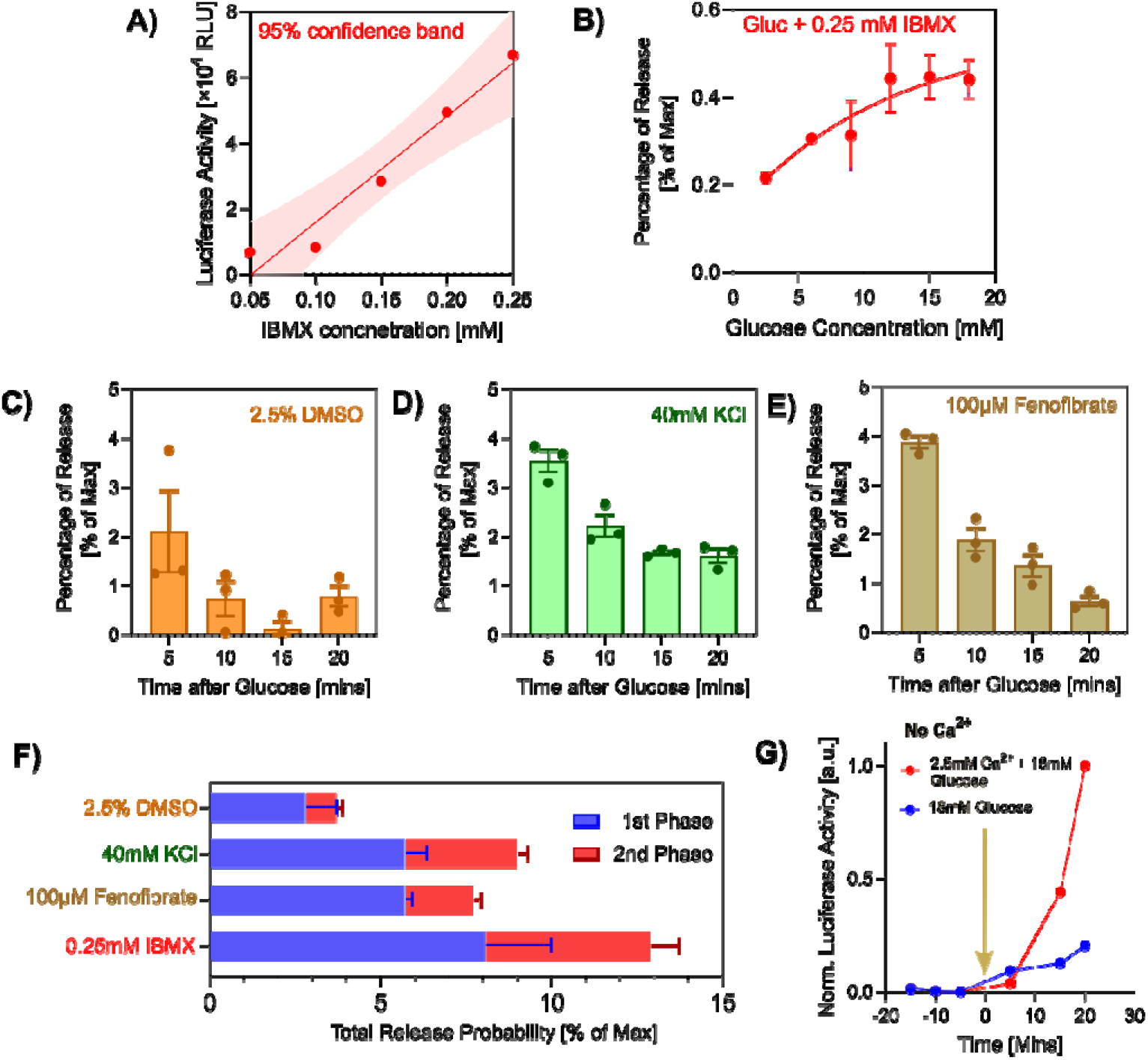
Quantitative assessment of Nluc secretion in biphasic modes with known secretagogues. **(A)** Dose-response relationship for IBMX (Isobutyl methylxanthine) at 18 mM glucose concentration in a stable cell line expressing Nluc; linear fit with 95% confidence band, R^2^=0.95. **(B)** The representative plot shows the percentage of Nluc released under conditions with and without 0.25 mM IBMX in 18 mM glucose; data represent mean ± SEM (n = 2). The representative bar chart shows Nluc secretion induced by known insulin secretagogues incubated with INS1 stable cell line for 20 mins with **(C)** DMSO, **(D)** potassium chloride (KCl), and **(E)** fenofibrate with 18 mM glucose; data represent mean ± SEM (n = 3). **(F)** Summary of the Nluc release probability in the first phase (blue) and the second phase (red) under known insulin secretagogues and suppressors; data represent mean ± SD (n = 3). (G) The chart shows the effect of glucose (blue) and extracellular Ca^2+^ (red) on Nluc secretion.

For the time series experiment, the supernatant was carefully removed from the wells without disturbing the cells. To minimize well-to-well variation in cell number, we ensured that the cells were firmly attached to the plate in order to evaluate different conditions in parallel. Additionally, we examined the effects of potassium chloride, dimethyl sulfoxide and fenofibrate on insulin secretion. When tested with potassium chloride compared to dimethyl sulfoxide (DMSO), a substance that depolarizes membranes and is widely used in vesicle fusion assays in live cells, insulin secretion was observed in both phases (Figure 2C & 2D). In our next investigation, we examined fenofibrate, a drug commonly used to treat hypertriglyceridemia, hypercholesterolemia, and dyslipidemia in cardiometabolic disease by lowering lipid levels in the liver and skeletal muscles.^30-32^ As expected, fenofibrate significantly inhibited luciferase secretion at higher glucose concentrations (Figure 2E) by acting as an agonist of the peroxisome proliferator-activated receptor α (PPARα) and facilitating fatty acid oxidation.^33^ Figure 2F shows the summary of the effect of known secretagogues and suppressors in beta cell secretion.

Next, we examined the relationship between free cytosolic Ca^2+^ concentration and insulin secretion by testing with and without extracellular Ca^2+^ in the buffer. The release of Nluc is only triggered in the presence of extracellular Ca^2+^ in the KRB (135 mM NaCl, 5mM KCl, 1 mM MgSO_4_, 0.4 mM K_2_HPO_4_, 5.5 mM Glucose, 20 mM HEPES) buffer, as shown in Figure 2G. Cosecretion of Nluc and Insulin is triggered when the membrane depolarizes, allowing calcium to enter rapidly into the cell. The amount of Ca^2+^ that enters voltage-dependent calcium channels could directly influence the rapid fast and progressive slow-release patterns in response to stimulation with KRB buffer. Overall, the platform opens a venue for dissecting the role of calcium signaling in beta cell secretion.

### 3.3. Genetic Screening Studies

In order to assess the suitability of the assay for genetic studies, VAMP8 knockdown and Syntaxin 4/SNAP23 overexpression were investigated, which are known to play an essential role in insulin secretion by regulating insulin-containing dense-core vesicles fusion to the plasma membrane. In addition, Syntaxin proteins may also be necessary for other activities, such as participating in endosomal recycling, cellularization during embryonic development, and GLUT4 translocation. With the aim of demonstrating that the results of our high-throughput assay based on rat INS 1 cells expressing Nluc are comparable with the results obtained with human islet perfusion assays, we performed VAMP8 gene knockdown studies. Western blotting results confirmed the absence of VAMP8 in the knockdown cells (Figure 3A). We examined the physiologic biphasic glucose-stimulated insulin secretion (GSIS) in VAMP8 knockdown cells using our assay. The results indicate that no reduction of luciferase signal in the first 5 mins indicating that predocked vesicles are not affected, but the releasable (newcomer) pool vesicles in the 10,15 and 20 mins are strongly affected (Figure 3B). As a result of inadequate recruitment of newcomers (releasable pool) vesicles to the plasma membrane, VAMP8 deficiency reduces IBMX-potentiated glucose stimulated insulin secretion (GSIS) by 84% in phase 1 and 87% in phase 2. (Figure 3C).

**Figure 3.**
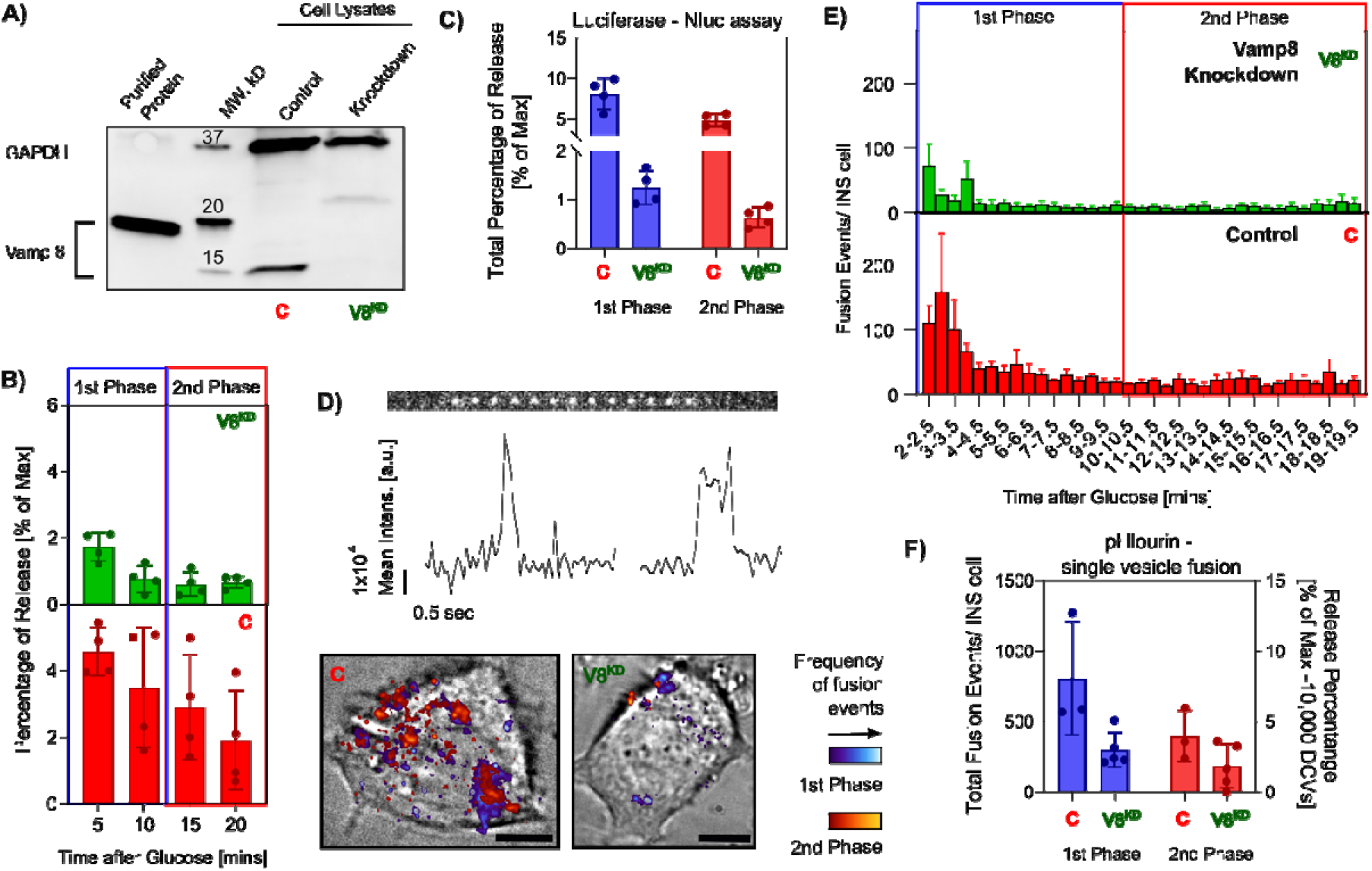
Quantitative assessment of the effect of VAMP8 deletion on insulin secretion in living cells using Nluc and single-vesicle fusion assays. **(A)** Western blotting analysis of VAMP8 knockdown (V8^KD^) INS 1 cells stably expressing Nluc. Purified VAMP8 protein and the control cells were used as controls. Purified VAMP8 protein is coupled with a 3x myc tag, therefore resulting in the size discrepancy. The efficiency of V8^KD^ is 91± 2.3%. **(B)** Histogram of the fusion events in the first phase and second phase in V8^KD^ (top) and control-C (bottom). Nluc assay; data represent mean ± SEM (n = 4). **(C)** Summary of the fusion events in the first phase (blue) and the second phase (red) for control and V8^KD^ from single vesicle fusion assay from Nluc assay. **(D)** Representative kymographs and fluorescence intensity curves. Fusion events are overlayed on the bright field, showing a decrease in vesicle fusion. **(E)** Histogram of the fusion events in the first phase and second phase in V8^KD^ and control (bottom). INS1 cells from single-vesicle fusion assay in the live cells; data represent mean ± SEM (n = 3 for control and n = 4 for V8^KD^). **(F)** Summary of the fusion events in the first phase (blue) and the second phase (red) for control and V8^KD^ from single vesicle fusion assay using proinsulin-pHlourin; data represent mean ± SEM (n = 3 for control and n = 4 for V8^KD^). Release percentage was computed with a maximum total dense core vesicle (DCV) count of ∼ 10,000 vesicles.

**Figure 4.**
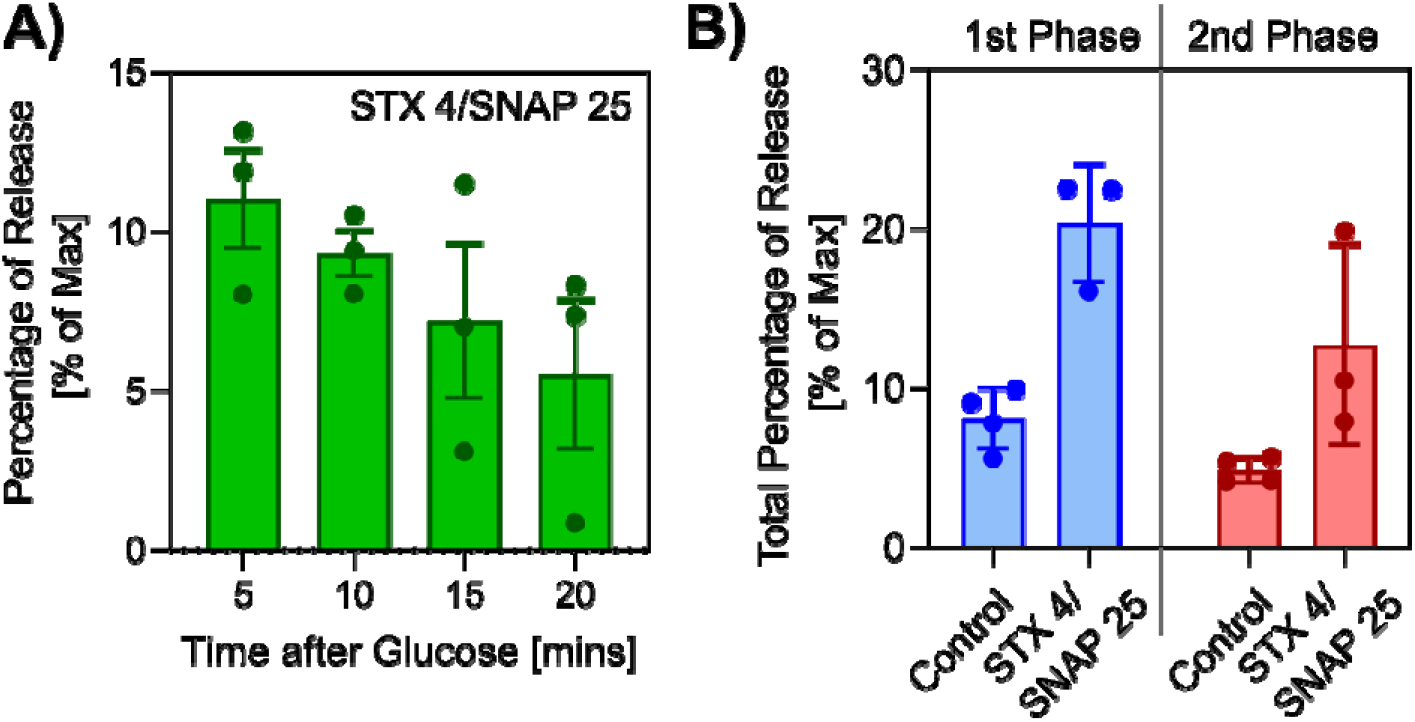
Effect of Syntaxin (STX) 4/SNAP23 overexpression on glucose-stimulated insulin secretion (GSIS) of pretreating INS-1E cells **(A)** The representative bar graphs indicates the release of luciferase at 5-minute intervals; data represent mean ± SEM (n = 3). **(B)** Summary of the luciferase secretion efficiency in the first phase (blue) and the second phase (red) for control and Syntaxin4/SNAP25 overexpression; data represent mean ± SEM (n = 4 for control and n = 3 for STX4/SNAP25).

Next, in order to investigate whether the Nluc and single-vesicle fusion assays performed in live cells are comparable, we used time-lapse total internal reflection fluorescence microscopy (TIRFM) to observe single cells for a period of 20 minutes to track exocytosis of insulin-dense core vesicles using Proinsulin-pHlourin. A pH-sensitive variation of green fluorescent protein called pHluorin is quenched by acidic environments (such as vesicle lumens) and becomes brighter in neutral environments (such as cell surfaces).^34^ As a result, true vesicle fusion events can be identified. A vector containing Proinsulin-pHlourin was transfected into the cells 24 hours before the experiment. At the basal unstimulated state, no measureable pHlourin is observed. Upon stimulating with 18 mM glucose and 0.25 mM IBMX, a bright flash was observed over the background signal indicating the release of insulin coupled with pHlourin (Figure 3D). Based on cumulative fusion events assessed over a 20-minute stimulation period, knockdown cells experienced lower cumulative fusion events than wild-type cells (Figure 3E). Syntaxin4 overexpression increases IBMX-potentiated glucose stimulated insulin secretion (GSIS) by 152% in phase 1 and 163% in phase 2.

A stable cell line expressing Nluc was transiently transfected once cells reached 60% confluency with a plasmid co-expressing Syntaxin4 and SNAP23. Western blot analysis of the transfected cell lysates showed an increased expression of these proteins compared to the cells without transfection. Overexpression of Syntaxin4 significantly increased insulin secretion in both the first and second phases.

### 3.4. Modeling disease environment

Beta cell toxicity has been associated with patients with Type 1 and Type 2 diabetes, along with defects in insulin secretion. To mimic these toxic conditions, previous studies have used cytokine treatment to simulate T1D pathology, leading to a reduction in insulin secretion due to the loss of DOC2B expression. Type 2 Diabetes was modeled using palmitate treatment, which resulted in a significant decrease in the first phase. In order to evaluate the assay for rapid screening of the effects of toxic environments on the actions of insulin biphasic secretion, saturated fatty acids, ER stress-inducing chemicals, and drugs were used to model beta-cell dysfunction.

Figure 5 shows the summary of two phases of insulin secretion dynamics affected by the external environment. First, we modeled a lipotoxic environment by treating cells expressing the luminescence reporter with sodium palmitate for 8 hours to induce lipotoxicity, since chronic exposure to free fatty acids impairs insulin secretion.^35,36^ We found a profound decrease in Nluc secretion when we stimulated the cells with higher glucose levels, a finding that is consistent with previous publications. Second, we tested the effect of ER stress on luciferase release using thapsigargin, a known SERCA inhibitor.^37-41^ In the presence of thapsigargin compound, glucose-stimulated Nluc secretion significantly decreased after 8 hours of treatment. Our results are consistent with previous investigations using isolated pancreatic islets showing that thapsigargin-induced ER stress alters protein folding and phosphorylation at Ser51 of the eukaryotic translation initiation factor (eIF2_α_).^42^ Cells can activate the unfolded protein response to restore protein folding balance and alleviate ER stress, thereby preventing cell death and restoring ER homeostasis. Finally, we tested the effect of the Streptozocin drug in our assay to model the type 1 diabetic pathology. The antibiotic streptozotocin destroys pancreatic islet cells and is commonly used as an experimental model for type 1 diabetes.^43-45^ As expected, Nluc secretion at high-glucose levels decreased when 10 mM streptozotocin was applied to the cells.

**Figure 5.**
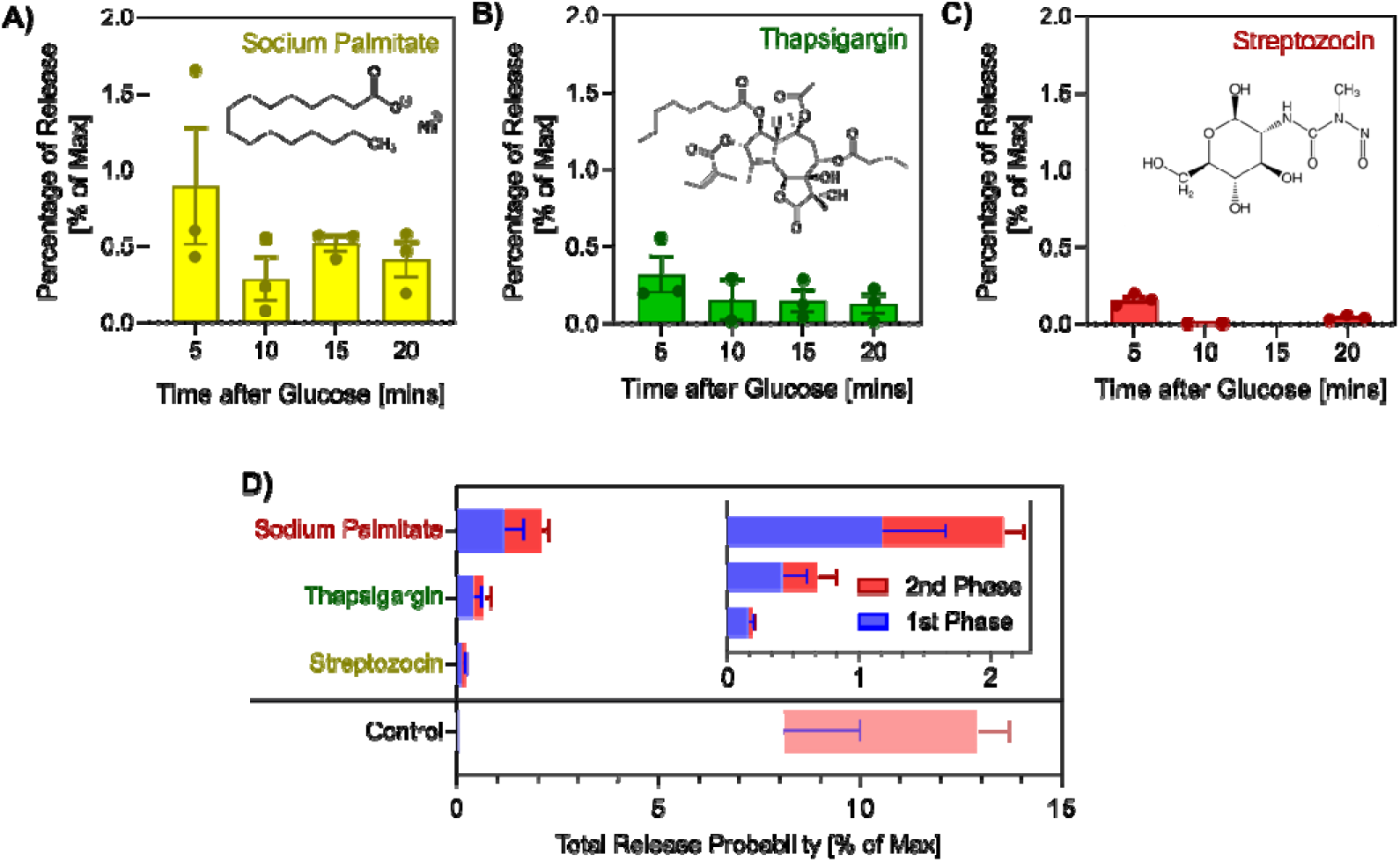
Effects of chemicals known to cause disease pathology on the first and second phase secretion in beta cells by GSIS **(A)** 8 hr pretreatment with sodium palmitate, saturated fatty acid is known to cause T1D disease pathology, suppresses insulin secretion. Data represents mean ± SEM (n = 3). **(B)** 8 hr pretreatment with thapsigargin, suppresses insulin secretion by causing ER stress. Data represents mean ± SEM (n = 3). **(C)** Treatment of streptozotocin affects the beta cells mass and a commonly used model for T1D pathology shows complete loss of insulin secretion. Data represents mean ± SEM (n = 3). **(D)** Results of a screen of three known drugs that are known to cause pathological states and affect the secretion in both the first phase (blue) and the second phase (red); data represent mean ± SD (n = 3).

## 4. Discussion

Reporter genes such as fluorescent proteins and luciferases that can be fused with the target of interest are useful for identifying the protein amounts, discovering signaling pathways, and monitoring protein-protein interactions. In this work, we have developed and validated a high-throughput assay based on a nanoluc luciferase to rapidly quantify the first and second-phase insulin secretion dynamics that facilitate large-scale assays of beta cell function. This is the first assay to quantify the percentage of glucose-mediated first and second phase insulin release at a specified time interval. The rapid nature, low cost, and high throughput of the Nluc assay makes it easier to identify molecular players involved in biphasic release. Since Nluc is small and monomeric, it is able to efficiently localize within insulin-containing secretory granules and co-secrete with insulin when high glucose levels are present. This method can be used as an alternative to ELISA, which is expensive and requires several liquid handling steps. A recent exploding growth in luciferase engineering will help us to perform a simultaneous assessment of multiple signaling pathways.^46-48^ We have demonstrated that the assay has the ability to provide a rapid and straightforward readout of insulin release during the first and second phases and the effect of beta-cell dysfunction on these phases. We also demonstrate that the assay is amenable to 96 well platform (SI Figure 6). The method thus paves the way for identifying the signaling pathways that affect insulin secretion as well as finding compounds that can reverse insulin secretion decline in diabetes through high-throughput compound screening.

To demonstrate the utility of the assay, cells were treated with chemicals and washed, then stimulated with glucose without treatment chemicals. Therefore, the suppression or enhancement of glucose-stimulated secretion can be related to changes in gene expression, protein levels, or post-transcriptional modifications. We establish the credibility of the assay with known secretagogues and suppressors that affect insulin secretion. We found that isobutyl-1-methylxanthine (IBMX) enhances the secretion in both first and second phase at results in increased luciferase secretion in the presence of 18 mM glucose. This indicates IBMX could be a perfect model compound together with glucose for studying the first and second-phase insulin secretion. Our assay on genetic screens tested the impact of knocking down the VAMP8 exocytosis protein resulting in no significant effects on pre-docked vesicles (first 5 minutes), but a significant impact on newcomer vesicles. The percentage of vesicle fusion is consistent with the results from single-vesicle exocytosis using proinsulin-pHlourin in live cells. Our overexpression Syntaxin4/SNAP23 studies suggest a strong increase in insulin secretion in both phases and could be a good target for type 2 diabetic therapy. Our screens also showed the impact of pathological agents on insulin’s first and second phase release. We will expand the screening platform in the future in order to screen all proteins that regulate exocytosis as well as genes that modulate lipid metabolism. By using a variety of treatment durations and phenotypic outcomes, we hope to uncover new pathways that are involved in glucose-regulated insulin secretion and ways to increase insulin secretion.

Since this assay focuses on Nluc secretion, the impact of insulin synthesis and insulin mutations are not addressed. We accept that is the limitation of the assay, but the Nluc is properly targeted, suggesting the proinsulin is properly made and loaded into the vesicles. We also accept that this method is very hard to quantify the fusion pore kinetics, amount of insulin release per vesicle, and single molecule protein imaging. Together with single-vesicle fusion assays,^49-55^ we can dissect the roles of individual molecular components in the insulin biphasic secretion process. This includes our patented suspended lipid membrane platform (SLIM),^56^ which uses synthetic reconstitutions or native lipid membranes to perform high-throughput fusion assays while also providing the precision to observe single molecule localization at the same time.

## 5. Conclusions

In summary, our team has created a quantitative assay with high throughput capability to analyze the first and second stages of insulin secretion. This assay enables the identification of beta cell signaling pathways responsible for regulating biphasic insulin secretion. Furthermore, the platform permits large compound libraries screening to discover substances that can suppress or enhance glucose-stimulated insulin secretion’s first or second phases. By precisely measuring luciferase secretion at regular time intervals and replicating the physiology, we’ve developed a cost-effective alternative to costly ELISA tests and human islet perfusion assays.

## Author contributions

ST, JW, MT, MB, JC, LL, MM, and SR performed the experiments. ST and JW authors analyzed the data, and SR wrote the manuscript.

## Acknowledgements

We thank Prof. James Rothman for the valuable discussions and for reading the manuscript. The authors thank Ramalingam Venkat Kalyana Sundaram for his help with experiments, valuable discussions, and for reading the manuscript. We also thank Sarvagya Saluja for her help with dense core vesicle isolation.

